# Plant Phenolics Inhibit Focal Adhesion Kinase and Suppress Host Cell Invasion by Uropathogenic *Escherichia coli*

**DOI:** 10.1101/2023.11.23.568486

**Authors:** Adam J. Lewis, Amanda C. Richards, Alejandra A. Mendez, Bijaya K. Dhakal, Tiffani A. Jones, Jamie L. Sundsbak, Danelle S. Eto, Matthew A. Mulvey

**Affiliations:** Division of Microbiology and Immunology, Department of Pathology, University of Utah, Salt Lake City, UT 84112, USA; School of Biological Sciences, 257 S 1400 E, University of Utah, Salt Lake City, UT 84112, USA; Henry Eyring Center for Cell & Genome Science, 1390 Presidents Circle, University of Utah, Salt Lake City, UT 84112, USA

## Abstract

Traditional folk treatments for the prevention and management of urinary tract infections (UTIs) and other infectious diseases often include plants and plant extracts that are rich in phenolic and polyphenolic compounds. These have been ascribed a variety of activities, including inhibition of bacterial interactions with host cells. Here we tested a panel of four well-studied phenolic compounds – caffeic acid phenethyl ester (CAPE), resveratrol, catechin, and epigallocatechin gallate – for effects on host cell adherence and invasion by uropathogenic *Escherichia coli* (UPEC). These bacteria, which are the leading cause of UTIs, can bind and subsequently invade bladder epithelial cells via an actin-dependent process. Intracellular UPEC reservoirs within the bladder are often protected from antibiotics and host defenses, and likely contribute to the development of chronic and recurrent infections. Using cell culture-based assays, we found that only resveratrol had a notable negative effect on UPEC adherence to bladder cells. However, both CAPE and resveratrol significantly inhibited UPEC entry into the host cells, coordinate with attenuated phosphorylation of the host actin regulator Focal Adhesion Kinase (FAK, or PTK2) and marked increases in the numbers of focal adhesion structures. We further show that the intravesical delivery of resveratrol inhibits UPEC infiltration of the bladder mucosa in a murine UTI model, and that resveratrol and CAPE can disrupt the ability of other invasive pathogens to enter host cells. Together, these results highlight the therapeutic potential of molecules like CAPE and resveratrol, which could be used to augment antibiotic treatments by restricting pathogen access to protective intracellular niches.

**IMPORTANCE:** Urinary tract infections (UTIs) are exceptionally common and increasingly difficult to treat due to the ongoing rise and spread of antibiotic resistant pathogens. Furthermore, the primary cause of UTIs, uropathogenic *Escherichia coli* (UPEC), can avoid antibiotic exposure and many host defenses by invading the epithelial cells that line the bladder surface. Here we identified two plant-derived phenolic compounds that disrupt activation of the host machinery needed for UPEC entry into bladder cells. One of these compounds (resveratrol) effectively inhibited UPEC invasion of the bladder mucosa in a mouse UTI model, and both phenolic compounds significantly reduced host cell entry by other invasive pathogens. These findings suggest that select phenolic compounds can be used to supplement existing antibacterial therapeutics by denying uropathogens shelter within host cells and tissues, and help explain some of the benefits attributed to traditional plant-based medicines.

## INTRODUCTION

Plants can produce thousands of phenolic compounds, which are defined as secondary metabolites comprised of at least one aromatic ring with one or more hydroxyl groups (1, 2). These diverse molecules can serve a variety of functions, which include the protection of plants from ultraviolet radiation, oxidative stress, herbivores, and microbial pathogens (2-4). The dietary consumption of plant phenolic compounds is linked with an array of health benefits ranging from anti-tumorigenesis to antimicrobial effects (2, 5-7). Especially intriguing are reports that phenolic and polyphenolic compounds derived from cranberry (*Vaccinium macrocarpon*) and other botanical sources may help protect against urinary tract infections (UTI) in some individuals (8-12). These infections, which are most often caused by strains of uropathogenic *Escherichia coli* (UPEC), are exceptionally common and prone to recur (13-16). About one-quarter of women will have at least one recurrent UTI (rUTI) within six months of a primary infection, and many individuals suffer multiple rUTIs per year (13, 17-21). The rampant dissemination and amplification of antibiotic resistant UPEC strains and other uropathogenic bacteria over the past two decades has greatly complicated the treatment of UTIs and stimulated widespread interest in alternate, supplemental therapies (11, 22-27).

There have been multiple clinical studies aimed at defining the effects of cranberry on UTI, but results have been mixed and difficult to compare due to heterogeneity in the types and quantities of cranberry products used, variations in study population characteristics, and disparate means of defining UTI (e.g. (28-31)). Despite these complications, recent systemic reviews and meta-analyses of published studies concluded that the consumption of cranberry products could significantly lower the risk of UTI in patients with a history of rUTIs (32-34). Oftentimes, bacteria that cause a rUTI are similar, or identical, to the bacteria that were responsible for the initial UTI (16, 21, 35-37). These and other observations suggest that environmental or in-host bacterial reservoirs may repetitively seed symptomatic UTIs in some people. Studies in mice and humans indicate the existence of UPEC reservoirs both within the gut and within the host cells that comprise the mucosal surfaces of the genitourinary tract (21, 37-42).

By using adhesive organelles known as type 1 pili to bind key host receptors, UPEC can trigger actin cytoskeletal rearrangements that promote the envelopment and internalization of bound bacteria (reviewed in (42)). Within bladder epithelial cells, bacteria that are not immediately expelled can either enter the cytosol and rapidly proliferate to form large but transitory intracellular bacterial communities, or the pathogens can establish small and seemingly quiescent, long-lived reservoirs within endosomal compartments (38, 43-47). Once in place, intracellular UPEC reservoirs are well-protected from host defenses and multiple frontline antibiotics, and are consequently difficult to eradicate (16, 38, 40, 42, 48-52). The inhibition of host cell invasion by UPEC could short-circuit cycles of rUTI that may be caused, in some individuals, by the repeated resurgence of intracellular bacterial reservoirs.

Several phenolic compounds derived from cranberry can inhibit UPEC adherence to host cells *in vitro*, but few have been examined for their effects on host cell invasion by uropathogenic bacteria (8, 11, 12, 53-55). A class of polyphenols known as proanthocyanidins (PACs), which are found in cranberry and many other plants, are well-studied inhibitors of UPEC adherence to host cells and can interfere with bacterial invasion of intestinal epithelial and Hela cells (56-62). Within the gut, PACs may inhibit host cell invasion by both inducing bacterial aggregation and by disrupting the actin cytoskeleton (60, 61). PACs may also impact UPEC colonization of the host via effects on bacterial stress response pathways, motility, biofilm development, iron metabolism, and toxin expression (55, 62-65). However, PACs likely have limited direct effects on either host cells or UPEC within the urinary tract, as these compounds are not well absorbed within the intestinal tract following consumption and are extensively metabolized by the gut microbiota (66-70). Some PAC-derived metabolites are absorbed within the gut and can later be detected in urine where *in vitro* assays suggest that they may protect against UTI by multiple mechanisms, including the inhibition bacterial adhesion to host cells (68). It is not yet clear if any of these PAC-derived metabolites can also impact bladder cell invasion independent of effects on bacterial adherence.

In this study, we probed the anti-invasion properties of four well-studied plant-derived phenolics: caffeic acid phenethyl ester (CAPE), resveratrol, catechin, and epigallocatechin gallate (EGCG). These phenolics are similar to many found in extracts from cranberry and a variety of other medicinal plants, and have been linked, at least tentatively, with protection against UTI (10, 66, 69, 71-76). Results presented here show that select phenolics can inhibit host cell invasion by UPEC, as well as other invasive pathogens. This inhibitory effect correlates with suppressed activation of Focal Adhesion Kinase (FAK), a key host regulator of F-actin dynamics.

## RESULTS

### CAPE and resveratrol inhibit host cell invasion by UPEC

The structures of CAPE, resveratrol, catechin, and EGCG, as well as representative sources of each of these phenolics, are shown in **Fig. 1A**. To examine potential effects of these compounds on UPEC-host cell interactions, we utilized standardized cell association and gentamicin protection invasion assays with the reference UPEC isolate UTI89 and the human bladder epithelial cell (BEC) line designated 5637 (77, 78). BECs were treated with each compound or carrier alone (DMSO) for 1 h prior to infection and maintained in the culture media throughout the 2-h cell association assays. During the course of these assays, the BEC monolayers remained alive and intact. None of the tested phenolic compounds altered the viability of UTI89 (**Fig. S1**), and only resveratrol caused a notable reduction in the numbers of cell-associated (intra- and extracellular) bacteria (**Fig. 1B**). In contrast, CAPE, resveratrol, and EGCG treatments significantly decreased the ability of UTI89 to invade the BECs relative to controls treated with only DMSO (**Fig. 1C**).

**Fig 1.**
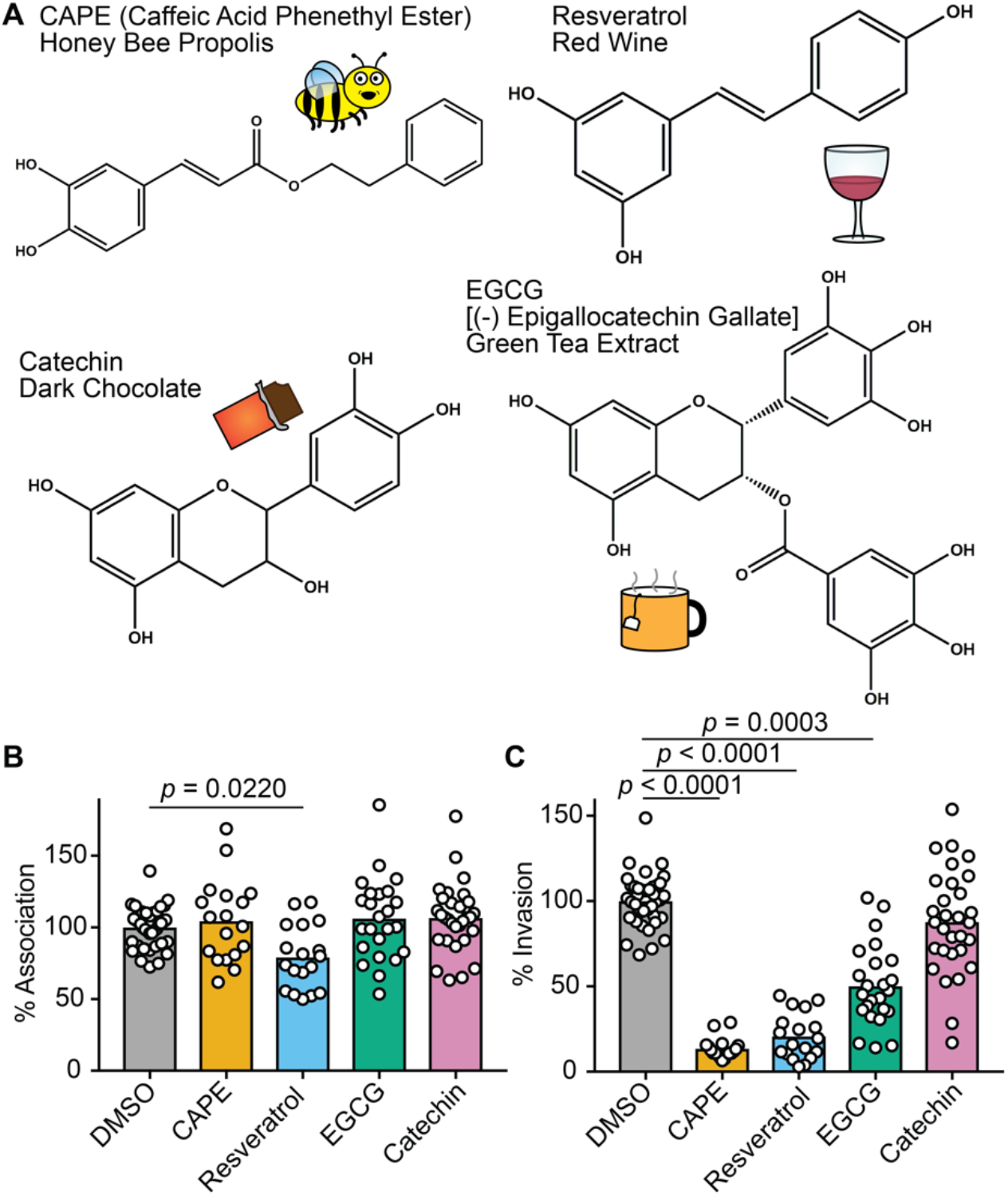
Phenolic compounds can inhibit host cell invasion by UPEC. (**A**) Skeletal structures of the phenolic compounds used in this study are depicted, with key dietary sources indicated via text and illustrations. (**B** and **C**) BECs were pretreated with CAPE (25 µg/mL), resveratrol (22.9 μg/mL), EGCG (25 µg/mL), catechin (25 µg/mL), or carrier alone (0.1% DMSO) for 1 h prior to infection with UTI89. Cells were then incubated for 2 h in the continued presence of the compounds, followed by a final 2-h incubation in medium containing gentamicin. Graphs indicate relative numbers of (**B**) cell-associated bacteria present prior to the addition of gentamicin and (**C**) intracellular, gentamicin-protected bacteria calculated as a fraction of the cell-associated bacteria. Data are normalized to DMSO-treated controls, with bars denoting mean values from at least three independent experiments performed in triplicate. *P* values were determined by Student’s *t* tests.

### BEC invasion by UPEC does not require de novo host transcription or translation

Previous studies indicated that CAPE, resveratrol, and ECGC can each inhibit activation of the host transcription factor NF-κB, which controls the expression of numerous genes, including many associated with inflammation and host responses to infection (79-81). With this information we reasoned that if the inhibitory effects of CAPE, resveratrol, and ECGC on UPEC invasion of BECs were attributable to the repression of NF-κB activation, then preventing downstream host transcriptional or translational responses processes should also interfere with UPEC entry into BECs. To test this idea, BECs were treated with actinomycin D (ActD) or cycloheximide (CHX), which ablate host transcription and translation, respectively (82). Neither drug impaired the ability of UTI89 to bind to or invade BECs (**Fig. 2**), indicating that the anti-invasion effects of CAPE, resveratrol, and ECGC are not due to the disruption of host transcription or translation downstream of NF-κB or other host transcription factors.

**Fig 2.**
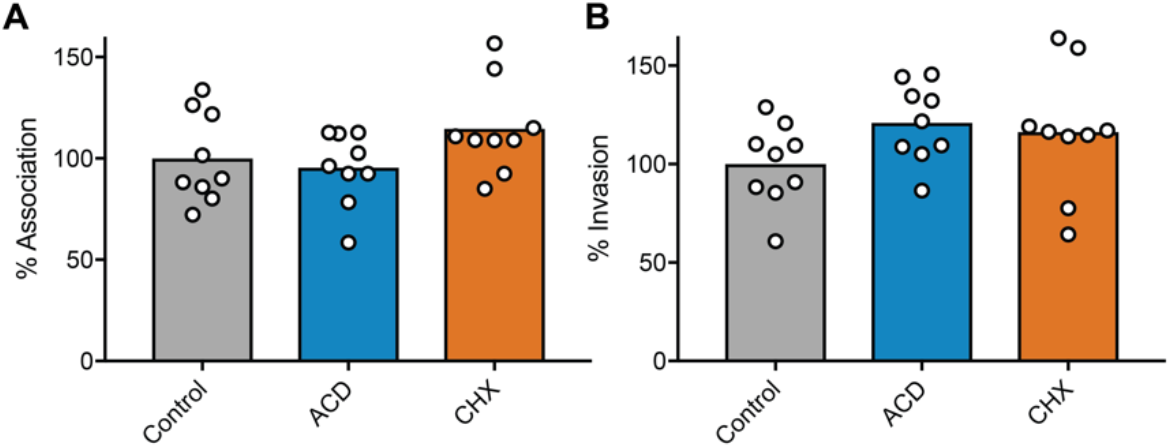
Host cell invasion by UPEC does not require active host transcription or protein synthesis. BECs were treated with actinomycin D (ACD, 5 µg/mL), cycloheximide (CHX, 26 µM), or carrier (ethanol) alone for 30 min and then infected in the continued presence of the inhibitors with UTI89 for 2 h followed by a 2-h incubation in medium containing gentamicin. Graphs show levels of (**A**) host cell-associated bacteria and (**B**) intracellular, gentamicin-protected bacteria, with bars indicating mean values. Data from three independent experiments performed in triplicate are expressed relative to controls that were treated with carrier (EtOH) alone. *P* values, as calculated by Student’s *t* tests, were all ≥ 0.28.

### CAPE and resveratrol inhibit FAK phosphorylation and increase focal adhesion numbers

Binding of the type 1 pilus-associated adhesin FimH to mannosylated glycoprotein receptors, including α3 and β1 integrins, activates host signaling cascades that drive the actin-dependent envelopment and internalization of bound UPEC (42). To examine phenolic effects on host signaling processes that might affect UPEC entry into BECs, we utilized an antibody microarray approach (Kinexus) to quantify changes in the phosphorylation of specific host proteins. For this assay we focused on CAPE, which had the greatest inhibitory effect on UPEC invasion (see **Fig. 1C**). Following a 15-min infection with UTI89, phosphorylated residues within several host factors that were previously linked with UPEC invasion were notably reduced (>25%) in CAPE-treated BECs relative to those that were treated with carrier alone (**Fig. 3A**) (42, 83-86). These factors included the FimH receptor β1 integrin, Akt (protein kinase B), vinculin, and FAK (protein tyrosine kinase 2), with phosphorylation of tyrosine 576 in FAK being the most diminished. Western blot analyses confirmed that CAPE treatment ablated FAK phosphorylation at Y576 [denoted as pFAK(Y576)] within UTI89-infected BECs, and showed that resveratrol had a similar effect (**Fig. 3B**). ECGC and catechin had less pronounced, but still discernable, effects on pFAK(Y576).

**Fig 3.**
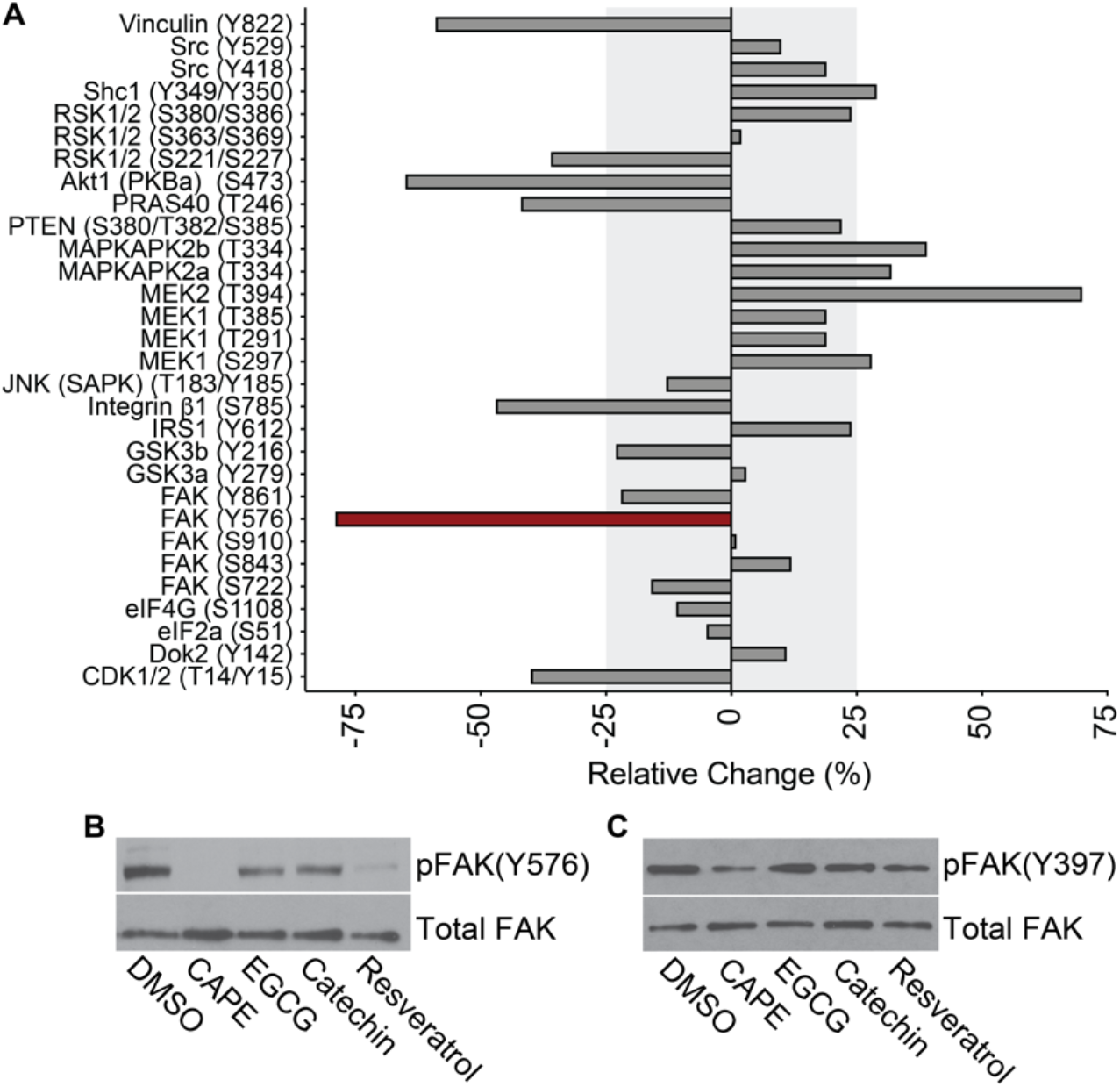
CAPE and resveratrol ablate phosphorylation of FAK at Y576. (**A**) Graph shows results from a Kinetworks Phospho-Site screen (KPSS 7.0), in which phosphorylation levels of each of the indicated residues (in parentheses) were quantified in CAPE- and DMSO-treated BECs after a 15-min infection with UTI89. Differences between samples are presented as percentages of the DMSO-treated, UTI89-infected controls: [(CAPE-treated – DMSO-treated)/DMSO-treated * 100]. Shaded areas denote relative changes of 25% or less, and the red bar highlights FAK(Y576) as the phospho-site most altered by CAPE treatment in this analysis (**B** and **C**) BECs were treated with carrier alone (0.1% DMSO), CAPE (25 µg/mL), EGCG (25 µg/mL), catechin (25 µg/mL), or resveratrol (22.9 μg/mL) for 1 h prior to a 15-min infection with UTI89 in the continued presence of each reagent. BEC lysates were then collected, resolved by SDS-PAGE, and probed by western blot analysis to assess (**B**) pFAK(Y576) and (**C**) pFAK(Y397) levels relative to total FAK in each sample.

FAK acts downstream of integrin receptors, working in concert with various signaling and scaffolding factors to modulate actin rearrangements and the maturation and turnover of focal adhesions (FAs) (87). These dynamic structures mediate actin-dependent host cell adherence and spreading processes, and a number of FA-associated factors, including FAK itself, are hijacked by UPEC and other pathogens to gain entry into host cells (42, 88). Integrin interactions with extracellular matrix proteins lead to the autophosphorylation of FAK at Y397, which in turn stimulates the recruitment and activation of SH2-domain-containing proteins such as phosphatidylinositol 3-kinase (PI3K) and Src kinase (87). Src then phosphorylates multiple sites within FAK, including Y576, which is required for maximal activation of FAK and the proper regulation of FA dynamics (87, 89). CAPE, but not the other phenolic compounds that we tested, diminished phosphorylation of FAK(Y397) within UTI89-infected BECs (**Fig. 3C**), but this effect was more subtle than what was observed with pFAK(Y576) (**Fig. 3B**).

In fibroblasts, the deletion of FAK increases the numbers of FA-like structures due to diminished turnover of integrin-linked adhesion sites (87, 90). By preventing full activation of FAK, we hypothesized that CAPE and resveratrol (and to a lesser extent EGCG and catechin) would partially mirror the effects of a FAK deletion and alter FA numbers. To test this possibility, uninfected BECs were treated with each phenolic compound individually or with carrier (DMSO) alone for 3 h and then processed for imaging by fluorescence confocal microscopy. Labeling of vinculin was used to visualize and quantify FAs, as previously described (91), and the BECs were counterstained to highlight nuclei and F-actin (representative images are shown in **Fig. 4A**). CAPE and resveratrol treatments both significantly increased the numbers FAs per cell (**Figs. 4A-B**) while EGCG and catechin slightly, but significantly, elevated the average size of the FAs (**Fig. 4C**). Together, these observations indicate that CAPE and resveratrol (more so than EGCG and catechin) can interfere with FAK activation and the turnover of FA-like complexes.

**Fig 4.**
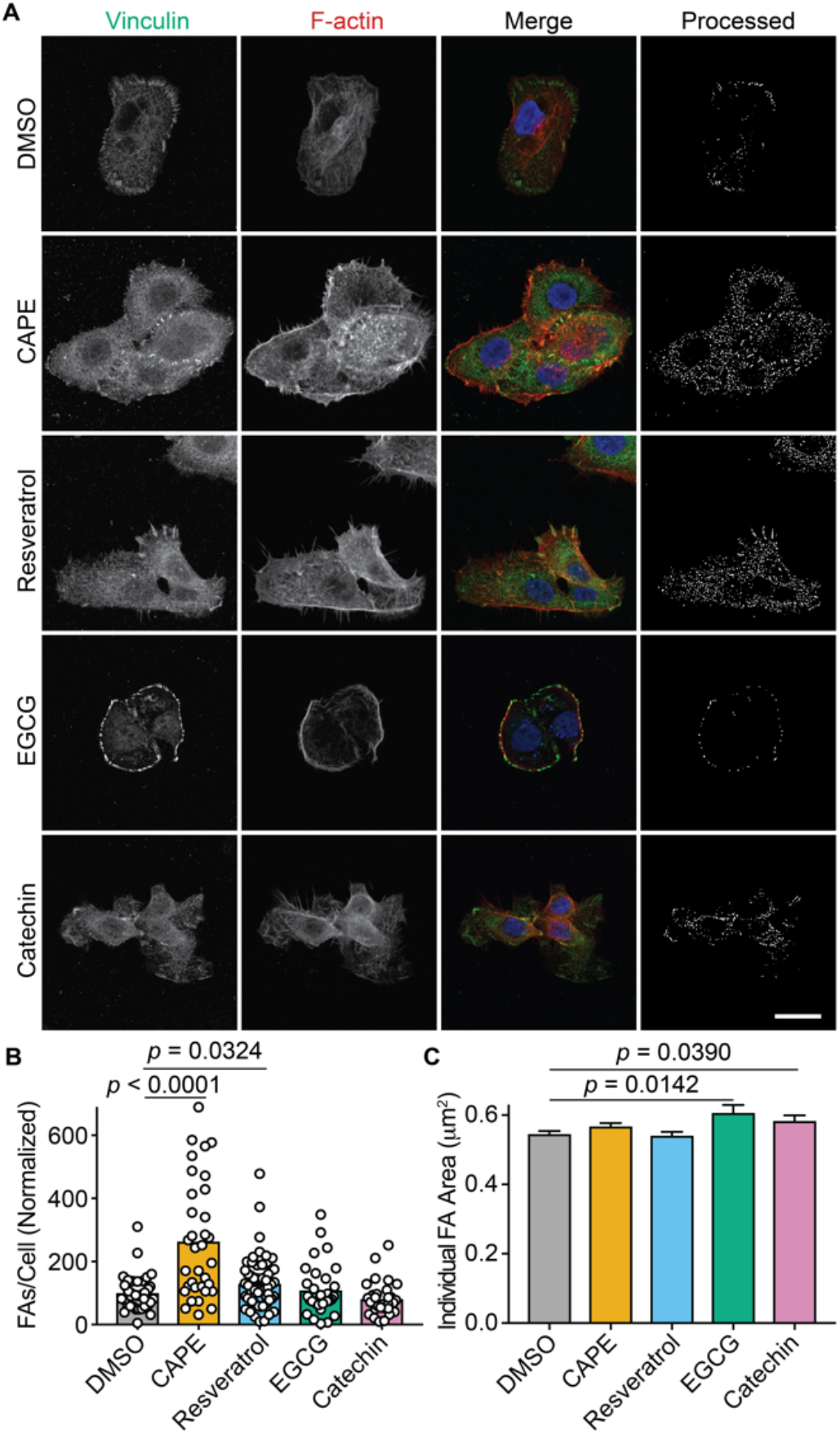
CAPE and resveratrol increase focal adhesion numbers. (**A**) Confocal microscopy images of BECs that were treated for 3 h with carrier alone (DMSO), CAPE (25 µg/mL), resveratrol (22.9 μg/mL), EGCG (25 µg/mL), or catechin (25 µg/mL), and then fixed and stained for vinculin (green), F-actin (red), and nuclei (blue). Single channel and merged images are indicated. The final panel in each row shows the cell images after processing to highlight focal adhesions for quantification. Scale bar, 10 μm. At least 30 cells from 3 independent experiments were processed to determine focal adhesion (**B**) numbers and (**C**) areas following the indicated treatments. Bars denote mean values (±SEM in C). *P* values were calculated relative to DMSO-treated controls by Student’s *t* tests.

### The anti-invasion effects of CAPE and resveratrol are largely attributable to FAK inhibition

Previously, the importance of FAK as a mediator of host cell invasion by UPEC was demonstrated using FAK-null (FAK-/-) mouse embryonic fibroblasts (MEFs) and siRNA with BECs (83). Building on this work, we treated BECs with a pharmacological inhibitor of FAK (FAK inhibitor 14, FAK14), which selectively blocks autophosphorylation of Y397 (92). We found that the treatment of BECs with FAK14 markedly reduced UTI89 internalization, but did not significantly alter the levels host cell-associated bacteria (**Fig. 5A-B**) nor bacterial viability in the culture medium (**Fig. S1**). These results echo those obtained using CAPE-, resveratrol-, and, to a lesser extent, EGCG-treated BECs (**Fig. 1B-C**).

**Fig 5.**
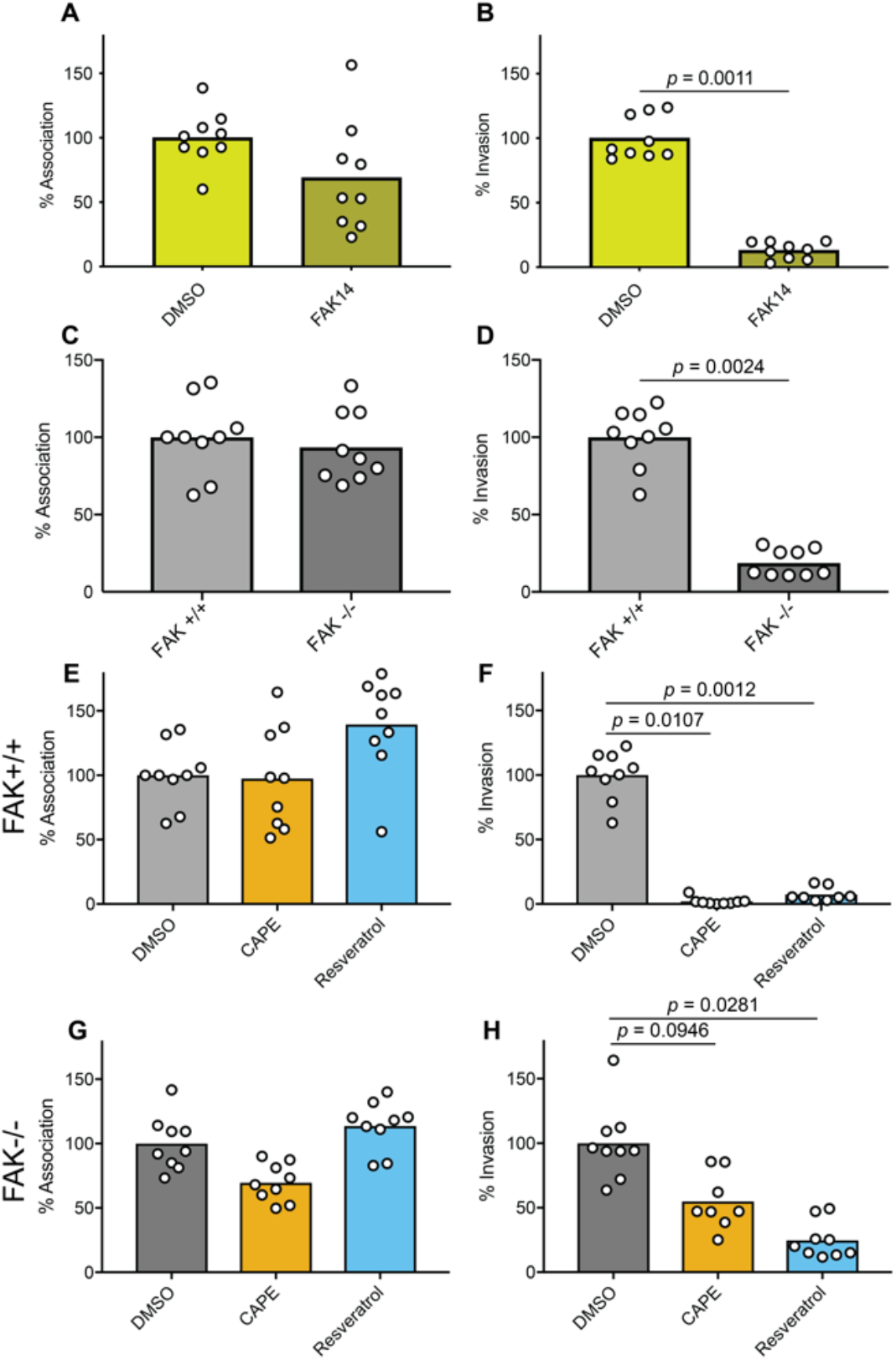
FAK inhibition and deletion mirror the effects of CAPE and resveratrol on host cell invasion by UPEC. (**A** and **B**) BECs or (**C-H**) FAK+/+ and FAK-/-MEFs were treated with FAK14 (10 μg/mL), CAPE (25 µg/mL), resveratrol (22.9 μg/mL), carrier (DMSO) alone, or left untreated, as indicated, for 1 h prior to and during a 2-h infection with UTI89. Monolayers were then washed and processed to determine total numbers of host cell-associated bacteria or incubated for an additional 2-h period with gentamicin to eradicate extracellular bacteria. Graphs show relative levels of (**A, C, E, G**) host cell-associated bacteria and (**B, D, F, H**) intracellular, gentamicin-protected bacteria. Data were normalized to DMSO-treated controls or to wild-type (FAK+/+) MEFs, as applicable, with bars representing mean values from 3 independent experiments carried out in triplicate. *P* values were determined by Student’s *t* tests.

Because CAPE and other phenolics can alter the phosphorylation patterns of multiple host factors (see **Fig. 3A**, e.g. (93-101)), we reasoned that the inhibitory effects of CAPE and resveratrol on UPEC invasion may not be entirely attributable to FAK inactivation. To address this possibility, we employed wild-type (FAK+/+) and FAK-null MEFs in combination with CAPE and resveratrol. As expected, UTI89 entry into the FAK-null MEFs was greatly impaired, though the bacteria bound the wild-type and FAK-/-host cells at similar levels (**Fig. 5C-D**). Treatment of the wild-type MEFs with either CAPE or resveratrol mirrored the effects seen with BECs (see **Fig. 1B-C**), suppressing host cell invasion by UTI89 while causing no significant changes in the total numbers of host cell-associated bacteria (**Fig. 5E-F**). Next, we asked if CAPE or resveratrol could also suppress UTI89 entry into FAK-null MEFs, which are already by and large refractory to host cell invasion by this pathogen (see **Fig. 1C**). Treatment of the FAK-null MEFs with CAPE led to somewhat reduced numbers of both bound and internalized bacteria, but these effects were not significant in comparison with DMSO-treated controls (**Fig. 5G-H**). In contrast, resveratrol significantly inhibited UTI89 entry into the FAK-null MEFs without altering the total numbers of bound bacteria. In total, these results indicate the ability of CAPE and resveratrol to obstruct host cell invasion by UPEC is mostly attributable to the inhibition of FAK. However, other as-yet undefined activities associated with these phenolics (and especially with resveratrol) can further suppress the invasion process independent of effects on FAK.

### CAPE and resveratrol inhibit host cell entry by distinct intracellular pathogens

Many microbial pathogens, in addition to UPEC, can invade non-phagocytic host cells via actin-dependent processes that are facilitated by FAK (88). To determine if CAPE, resveratrol, EGCG, or catechin affect host cell entry by other invasive pathogens, we employed our standard cell association and invasion assays with *Salmonella enterica* serovar Typhimurium, *Shigella flexneri*, and a non-pathogenic surrogate for *Yersinia pseudotuberculosis* (AAEC185/pRI203). The latter is a type 1 pilus-negative K-12 *E. coli* strain that expresses the *Y. pseudotuberculosis* invasin protein, which promotes actin- and FAK-dependent host cell entry by binding integrin receptors (102, 103). None of the tested phenolics significantly altered the numbers of host cell-associated AAEC185/pRI203, though the numbers of adherent bacteria recovered from EGCG- and catechin-treated host cells trended higher and had a greater spread (**Fig. 6A**). As seen with UTI89, CAPE, resveratrol, and, to a lesser extent, EGCG significantly impeded host cell invasion by AAEC185/pRI203 (**Fig. 6B**). Similar results were obtained with *S. flexneri* (**Fig. 6C-D**), which mobilizes multiple type III secretion system effectors that engage integrin receptors and associated host factors to promote FAK phosphorylation coordinate with actin rearrangements that drive bacterial internalization (104).

*S.* Typhimurium can also use type III effectors to enter host cells via FAK- and actin-dependent processes, but the *Salmonella* effectors are distinct from those encoded by *Shigella* (105). Furthermore, though *S.* Typhimurium entry into host cells requires FAK, the kinase domain which contains the activating phosphosite Y576 is dispensable for host cell invasion by this pathogen (105). In our assays, none of the tested phenolics altered the levels of host cell-bound *S.* Typhimurium (**Fig. 6E**), and only resveratrol inhibited host cell invasion (**Fig. 6F**). Together, these findings indicate that the ability of CAPE, resveratrol, and EGCG to impede host cell invasion can vary markedly, dependent on the pathogen and its specific mechanism of entry.

**Fig 6.**
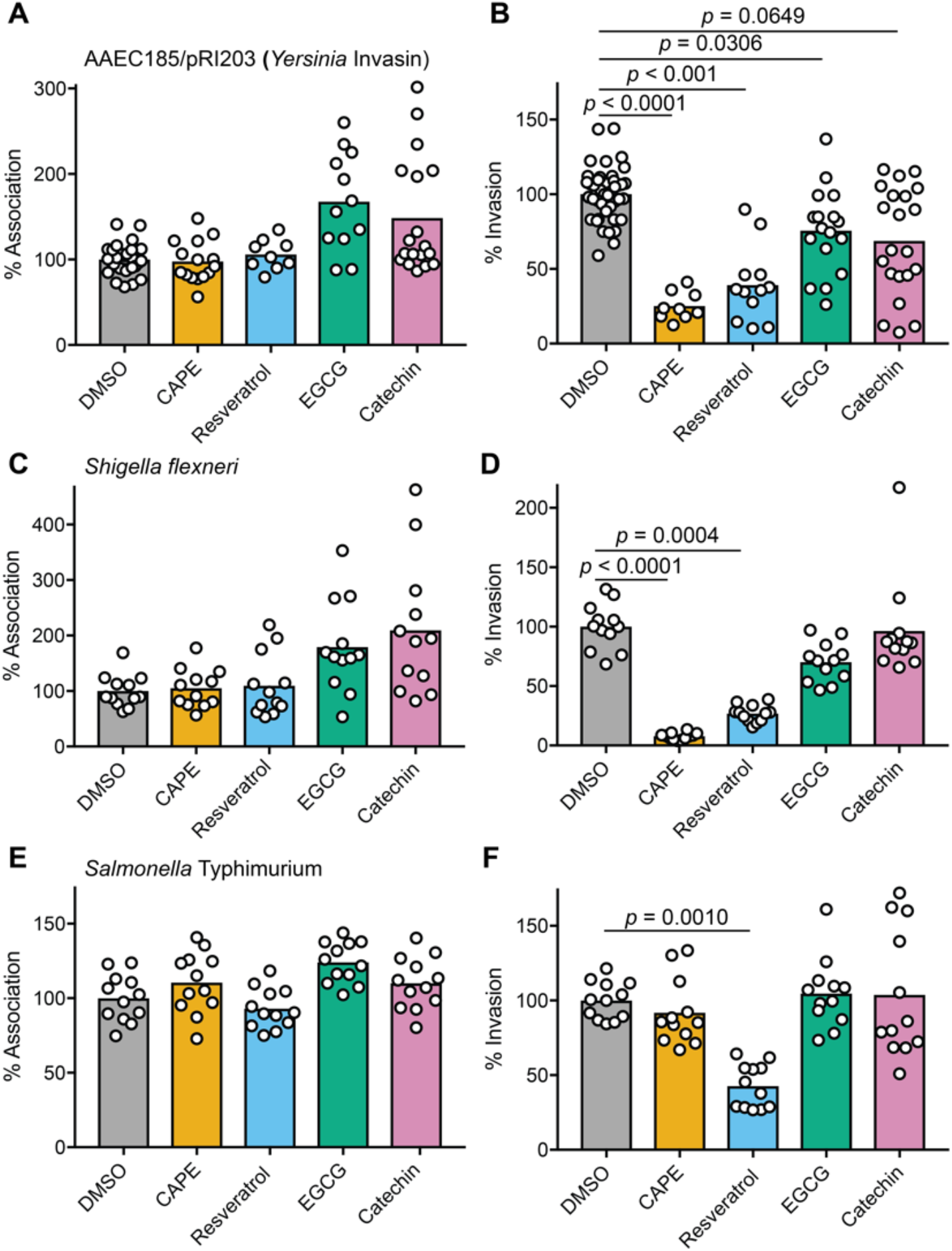
CAPE and resveratrol can inhibit host cell entry by other invasive bacteria. BECs were treated with the indicated phenolic compounds or DMSO alone for 1 h prior to infection with (**A** and **B**) AAEC185/pRI203, (**C** and **D**) *S. flexneri* (**E** and **F**), or *S.* Typhimurium. Monolayers were then incubated for 2 h in the continued presence of the compounds, followed by a 2-h incubation in the presence of gentamicin. Graphs show mean values of (A, C, E) cell-associated and (B, D, F) gentamicin-protected, intracellular bacteria from at least 3 independent experiments performed in triplicate. Data are expressed relative to DMSO-treated controls. *P* values were calculated using Student’s *t* tests.

### Resveratrol inhibits UPEC invasion of the murine bladder mucosa

Next, we tested if resveratrol could interfere with UPEC invasion of host cells in an established mouse model of UTI (41, 51). For this initial *in vivo* work, we focused on resveratrol as it was much more soluble than CAPE in both DMSO and in aqueous solutions, and consequently less prone to precipitate out when introduced into the bladder (106). Adult female CBA/JCrHsd mice were inoculated via transurethral catheterization with ∼10^7^ CFU of UTI89 in PBS containing 300 µM resveratrol or just the carrier DMSO. After 1 h, the bladders were collected, rinsed, and treated with gentamicin to kill any extracellular bacteria. Over 25-fold fewer intracellular bacteria were recovered from the resveratrol-treated bladders relative to those treated with DMSO alone (**Fig. 7**). These results indicate that resveratrol has the capacity to effectively inhibit host cell invasion by UPEC within the murine bladder.

**Fig 7.**
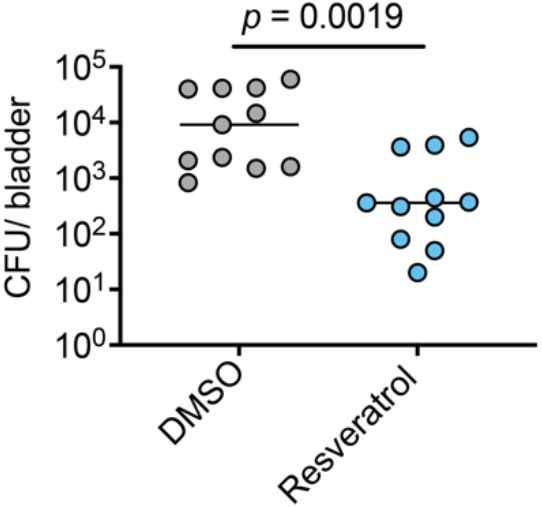
Resveratrol inhibits bacterial invasion of the bladder mucosa. Adult female CBA/JCrHsd mice were inoculated via trans-urethral catheterization with 10^7^ CFU of UTI89 suspended in PBS containing either DMSO or resveratrol (300 µM). Bladders were extracted 1 h later and the numbers of intracellular, gentamicin-protected bacteria were determined. Bars indicate median values; n=11 mice from two independent experiments. *P* value determined using the Mann-Whitney U test.

## DISCUSSION

Results presented here show that the plant phenolics CAPE, resveratrol, and, to a lesser extent, EGCG can inhibit UPEC entry into BECs. These phenolics are similar to compounds derived from cranberry-associated PACs and other complex polyphenolic biomolecules like tannins, which are attributed with a variety of antimicrobial activities including bactericidal and anti-adhesion effects (55, 58, 72, 107-109). In our assays, none of the tested phenolics interfered with bacterial viability (**Fig. S1**), and only resveratrol had a noticeable (though slight) inhibitory effect on UPEC adherence to BECs (**Fig. 1B**). Furthermore, we found that host cell invasion by UPEC did not require *de novo* host transcription or translation, indicating that the inhibitory effects of CAPE, resveratrol, and EGCG are not related to the ability of these phenolics to interfere with host transcription factors like NF-κB (**Fig. 2**). Rather, the more pronounced inhibitory effects of CAPE and resveratrol on UPEC entry into BECs were linked with the dysregulation of host actin dynamics via the suppression of FAK phosphorylation at Y576. The disruption of FAK signaling appears to be an effect of many plant-derived phenolic compounds (e.g. curcumin, enterolactone, glabridin (99-101, 110-114)), and may help explain some of the reported benefits of these molecules for the prevention or treatment of infections, cancers, and other ailments.

Extracts from a variety of medicinal plants, including *Citrus reticulata* Blanco (mandarin seeds), *Amaranthus caudatus* (a flowering plant that thrives in temperate-tropical areas), *Clinopodium bolivianum* (an aromatic shrub from the Andes region of South America), and *Lactuca indica* (Vietnamese dandelion) have been shown to inhibit both UPEC adherence to and invasion of bladder cells *in vitro* (115-118). Like cranberry, these plants are rich in phenolic compounds (117, 119-122), but the specific extract components that inhibit UPEC entry into BECs were not defined. Mechanistically, these extracts did not have any direct antibacterial activities and instead appeared to interfere with the invasion process by downregulating host cell receptors for type 1 pili or by suppressing downstream cell signaling events (115-118).

In the case of *L. indica* extract, the inhibition of BEC invasion by UPEC was partially attributable to the attenuation of FAK phosphorylation at Y397 (118). The autophosphorylation of this site, as noted above, is a proximal step leading to the recruitment of other signaling factors like PI3K and Src kinase that precede full activation of FAK and the instigation of FAK-modulated actin cytoskeletal rearrangements (87). In our assays, the effects of CAPE and resveratrol on the phosphorylation of FAK(Y397) were much less pronounced than those observed with FAK(Y576) (see **Fig. 3**), suggesting that these two phenolic compounds act further downstream in the FAK activation pathway than *L. indica* extract. Our experiments with FAK-null cells confirm that FAK is a major, though likely not the sole, host cell target that explains the inhibitory effects of CAPE and resveratrol on BEC invasion by UPEC (**Fig. 5**). This conclusion is corroborated by observations showing that CAPE and resveratrol treatments both lead to marked increases in the numbers of FAs, coordinate with alterations in the actin cytoskeleton (see **Fig. 4**).

Our observations with *S.* Typhimurium, *S. flexneri*, and recombinant *E. coli* expressing the invasin protein from *Y. pseudotuberculosis* reveal that the anti-invasion effects of CAPE and, especially, resveratrol can extend beyond UPEC (see **Fig. 6**). Of note, FAK can modulate host cell entry by each of these microbes (102, 104, 105). However, the differential effects of CAPE and resveratrol on host cell entry by a pathogen like *S.* Typhimurium (see **Fig. 6F**) suggest that these phenolics can have additional, non-overlapping effects on host cell processes that promote invasion, independent of FAK. This possibility is supported by multiple reports indicating that both CAPE and resveratrol can disrupt various host factors and signaling cascades that might directly or indirectly impact host cell invasion and intracellular trafficking by bacterial pathogens (e.g. (123-128)). The potential of plant-derived phenolic compounds to interfere with host cell invasion independent of effects on FAK is exemplified by luteolin, a secondary polyphenolic metabolite that is found in many fruits, vegetables, and medicinal herbs (129). Luteolin can limit UPEC entry into BECs by inhibiting host cAMP-phosphodiesterases, which in turn interferes with actin rearrangements driven by the activation of Rac1 GTPase.

There is growing interest in the development of therapeutics that can ameliorate disease by targeting host factors that are highjacked by microbial pathogens rather than the pathogens themselves (130-132). If effective, such host-directed therapeutics are expected to help sidestep the growing challenge of antibiotic resistance. Results with resveratrol-treated mice (**Fig. 7**) indicate that this phenolic, or compounds with similar activities, could be valuable therapeutic options that can deny UPEC refuge within host cells. Without the ability to hide within host cells, UPEC would be more susceptible to clearance by host defenses and antibiotic treatments that are often ineffective against intracellular microbes (16, 50). Phenolic compounds derived from cranberry, if able to act within the urinary tract in a similar fashion to resveratrol, could help explain the potentially beneficial effects of consuming cranberry products by some individuals who suffer from rUTI (32-34). The benefits of such phenolics could vary dependent on the cause, or source, of the rUTIs. These recurrent infections may arise via repeated inoculation of the urinary tract by pathogens acquired from environmental sources, from bacterial reservoirs within the gut, or from the resurgence of intracellular populations that can persist within the vaginal or bladder mucosa (21, 37-40). We speculate that inhibitors of UPEC invasion like CAPE and resveratrol might primarily aid the latter group, by interrupting cycles of intracellular persistence, growth, resurgence, and re-invasion of host cells within the genitourinary tract.

Though the use of resveratrol, CAPE, or functionally homologous compounds from cranberry or other sources as a means to combat UTI is an appealing notion, it should be tempered with an appreciation of the many obstacles associated with such an approach. Instillation of phenolic compounds by intravesical catheterization is not facile nor cost-effective, and agents delivered in this manner may not remain soluble or might not effectively penetrate target host cells within the mucosa. Furthermore, timing of this treatment approach may be complicated if the compounds need to be present prior to invasion, or re-invasion, of the mucosa by UPEC. Oral administration of anti-invasion phenolic compounds faces similar challenges, in addition to potential problems with absorption and modification by metabolic processes and microbes within the gut (53, 69, 133-135). Furthermore, the intake of very high amounts of plant-derived phenolics might have detrimental effects, such as iron depletion, liver and kidney toxicity, and irritation of the gastrointestinal tract (10, 12, 136-138). Despite these limitations, the work presented here highlights the potential therapeutic utility of plant-derived phenolic compounds as a means to inhibit host cell invasion by UPEC which, if optimized, could help disrupt cycles of rUTIs in some individuals.

## MATERIALS AND METHODS

### Bacterial strains, cell culture, and inhibitors

The UPEC cystitis isolate UTI89 was grown statically from frozen stocks for 24 h at 37°C in either LB (Difco) or modified M9 minimal medium to induce expression of type 1 pili (6 g L^-1^ Na2HPO4, 3 g L^-1^ KH2PO4, 0.5 g L^-1^ NaCl, 0.1 m*M* CaCl2, 1 g L^-1^ NH4Cl, 1 m*M* MgSO4, 0.1% Glucose, 0.0025% nicotinic acid, 0.2% casein amino acids, and 16.5 µg mL^-1^ thiamine) (23, 38). *S.* Typhimuirum (SL1344) and *S. flexneri* (ATCC 12022) were grown shaking at 37°C in LB overnight, then diluted 1:33 in fresh LB and grown for an additional 3.5 h, as previously described (139). AAEC185/pRI203 was grown shaking in LB to stationary phase prior to use (78, 140).

The bladder carcinoma cell line 5637 (ATCC HTB-9) was grown and maintained in RPMI1640 (HyClone) supplemented with 10% heat inactivated fetal bovine serum (FBS; HyClone) in a 37°C humidified incubator with 5% CO2. FAK +/+ (ATCC CRL-2645) and FAK -/- (ATCC CRL-2655) MEFs were grown and maintained in DMEM (HyClone) supplemented with 10% heat inactivated FBS.

FAK14 (a.k.a. Y15) was purchased from Cayman Chemical, while CAPE, resveratrol, EGCG, and catechin were from Sigma-Aldrich, Biomol, or Cayman Chemical. These compounds were prepared as 1000X stocks in DMSO. Actinomycin D and cycloheximide were obtained from Sigma-Aldrich and solubilized in ethanol.

### Bacterial cell association and invasion assays

Bacterial host cell association and invasion assays were performed using established protocols (78, 141). Briefly, 5637 or MEF cells were seeded into 24-well tissue culture plates and grown for about 24 h to near confluency. Where indicated, cell monolayers were treated with CAPE (100.7 μM; 25 µg/mL), resveratrol (100 µM; 22.9 μg/mL), EGCG (54.5 μM; 25 µg/mL), catechin (86.1 μM; 25 µg/mL), FAK14 (10 μg/mL; 35 μM), or DMSO (carrier, 0.1% final concentration) in complete media for 1 h prior to infection. Alternatively, host cells were treated with actinomycin D (5 µg/mL), cycloheximide (26 µM), or an equal volume of ethanol (diluent) for 30 min prior to infection. Triplicate sets of host cells were then infected with UTI89 or AAEC185/pRI203 using an MOI of approximately 15, while an MOI of 100 was used with *S. flexneri* and *S.* Typhimurium. Plates were centrifuged at 600 X g for 5 min to accelerate and synchronize bacterial contact with the host cells. UTI89- and AAEC185/pRI203-infected monolayers were then incubated at 37°C in the continued presence of the compounds or carrier, washed 4 times with PBS containing Mg^+2^ and Ca^+2^ (PBS^+2^), and lysed in PBS with 0.4% Triton-X 100. Serial dilutions of these lysates were plated on LB agar to determine numbers of cell-associated bacteria. Alternatively, sets of triplicate wells were washed twice with PBS^+2^ and treated for 2 h with complete media containing gentamicin (100 µg/mL) to kill extracellular bacteria. Subsequently, monolayers were washed 4 times with PBS^+2^ and lysed and plated as noted above to quantify the numbers of surviving intracellular bacteria. Experiments with *S.* Typhimurium and *S. flexneri* used 30-min infection periods for the cell association assays, followed by 1-h incubations with gentamicin for the invasion assays. Results from the invasion assays were normalized by dividing the numbers of intracellular bacteria by the total number of cell-associated bacteria, accounting for any differences in host cell numbers. All assays were repeated at least 3 times in triplicate.

Potential effects of the phenolic compounds and FAK14 on bacterial growth and viability were assessed by adding bacteria to complete RPMI media in 24-well plates using the same times and drug concentrations as used for the cell association assays, but in the absence of host cells. Bacterial titers were then determined by plating serial dilutions of the media on LB agar. These assays were independently repeated 3 times.

### Signal Transduction Protein Phospho-site Profiling

Sub-confluent 5637 BEC monolayers in 6-well plates were treated with CAPE (25 µg/mL) or DSMO alone for 1 h, infected with UTI89 (MOI∼15), and centrifuged at 600 X g for 5 min. After an additional 15 min incubation at 37°C in the continued presence of CAPE or DSMO, wells were washed 3 times with PBS^+2^, then lysed on ice with cold buffer containing 50 mM Tris (pH 7.4), 1 mM NaCl, 1% NP-40, complete protease inhibitor cocktail (Roche Applied Science), 1 mM PMSF, 1 mM NaF, 0.4 mM orthovanadate, 5 µM leupeptin, and 1 mM aprotinin. Protein concentrations were determined using a BCA reagent system (Pierce). Lysates were diluted in 4X Kinexus sample buffer to a final concentration of 0.8 µg/µL and shipped to Kinexus (Vancouver, Canada) for multi-immunoblotting analysis using the Kinetworks signal transduction protein phospho-site profiling service (KPSS 7.0 Profile).

### Western blot analysis

Nearly confluent BEC monolayers grown in 12-well plates were serum starved overnight, treated with the specified phenolic compounds or 0.1% DMSO alone for 1 h, and infected with UTI89 from M9 cultures using an MOI of about 25. The cell culture media was not exchanged when adding either the compounds or during the infection process. After a 5-min spin at 600 X g, the plates were incubated for 15 min at 37°C, washed 3 times with PBS^+2^, and then lysed in ice-cold RIPA buffer supplemented complete protease inhibitor cocktail (Roche Applied Science), 1 mM PMSF, 1 mM NaF, and 0.4 mM orthovanadate. Equivalent protein amounts (as determined by BCA assays; Pierce) were resolved by SDS-PAGE, transferred to Immobilon PVDF-FL membrane (Millipore), and processed for western blot analysis using mouse anti-FAK antibody (BD Biosciences) or phosphosite-specific mouse anti-pFAK(Y397) (BD Biosciences) and rabbit anti-pFAK (Y576) (Upstate Biotechnology) primary antibodies, and horseradish peroxidase-conjugated secondary antibodies (78, 141, 142).

### Visualization and Quantification of Focal Adhesions

5637 BECs were seeded onto 12 mm diameter coverslips in 24-well plates and grown overnight until nearly confluent. Cells were treated with CAPE (25 µg/mL), resveratrol (22.9 μg/mL), EGCG (25 µg/mL), catechin (25 µg/mL), or DMSO (carrier, 0.1%) alone in complete RPMI media for 3 h, washed 3 times with PBS^+2^, and then fixed for 20 min with 3.7% paraformaldehyde dissolved in PBS. After 3 washes in PBS, cells were blocked and permeabilized using PBS containing 1% powered milk, 3% bovine serum albumin, and 0.1% saponin. The cells were then labeled using primary mouse anti-vinculin antibody (1:100; Sigma-Aldrich) and donkey anti-mouse Alexa Fluor 555-conjugated secondary antibody (1:400; Abcam). F-actin and nuclei were stained using Oregon Green 488-conjugated phalloidin (1:200; ThermoFisher) and Hoechst (1:1000; Sigma-Aldrich), respectively. Coverslips were mounted in FluorSave (Calbiochem) and imaged using a Nikon A1 series confocal microscope with NIS Elements software.

Quantitative analysis of vinculin-positive focal adhesions was performed as previously described, with slight modifications (91). Briefly, using the Fiji processing package with ImageJ software the background for each image of vinculin-stained cells was subtracted and local contrast enhanced using the CLAHE plugin (143). Next, a mathematical exponential was utilized via the Exp function to further reduce background, and brightness and contrast were adjusted automatically. A Gaussian filter was applied using the Log3D plugin with sigma X=1.5 and sigma Y=1.5. An automatic threshold function was then used to create binary images in which pixels were assigned to either a background or foreground signal. Particles (representing focal adhesions) within the binary images were enumerated and sized using the ANALYZE PARTICLES command in ImageJ, with the size parameter set at 14.5-infinity.

### Mouse Infections

Using established protocols approved by the University of Utah and Institutional Animal Care and Use Committee (IACUC), 8 to 9-week-old female CBA/JCrHsd mice (Harlan Laboratories) were inoculated via transurethral catheterization with 10^7^ CFU of UTI89 in 50 µL PBS containing 300 µM resveratrol or DMSO (144). Mice were sacrificed 1 h post-catheterization and the bladders were harvested aseptically, quadrisected, and incubated for 30 min at 37°C in PBS with gentamicin (100 µg/mL) to kill extracellular bacteria. The bladder pieces were then washed 3 times with PBS and homogenized in PBS containing 0.025% Triton X-100. Serial dilutions of each homogenate were plated on LB agar to determine numbers of intracellular bacteria. A total of 11 mice from two independent experiments were tested for each treatment.

### Statistics

For the mouse experiments, data distribution normality (Gaussian) was not assumed. Mann–Whitney *U* tests and unpaired two-tailed Student’s *t* tests were performed using Prism 9.0.0 (GraphPad Software). *P* values of less than or equal to 0.05 were considered significant.

## ACKNOWLEDGEMENTS

This work was funded in part by NIH grants GM134331, AI095647, and DK069526 to MAM. Additional support was provided by NIH Microbial Pathogenesis T32 training grant AI055434 to ACR and DSE, NIH Hematology T32 training grant DK7115 to AJL, and a University of Utah Chevron Undergraduate Research Scholarship for women in STEM to AAM. The funders had no role in study design, data collection and interpretation, or the decision to submit the work for publication. The authors declare that there are no competing interests.

## CONTRIBUTIONS

Conceived and designed the studies: M.A.M., A.C.R., A.J.L., B.J.K., and D.S.E. Collected the data: A.C.R., A.J.L., A.A.M, B.J.K., D.S.E., T.A.J., J.L.S., and M.A.M. Data analysis: A.C.R., A.J.L., M.A.M., and B.J.K. Created the figures: A.C.R. and M.A.M. Wrote the paper: M.A.M., A.C.R, and A.J.L.

## Figures

**Fig S1.**
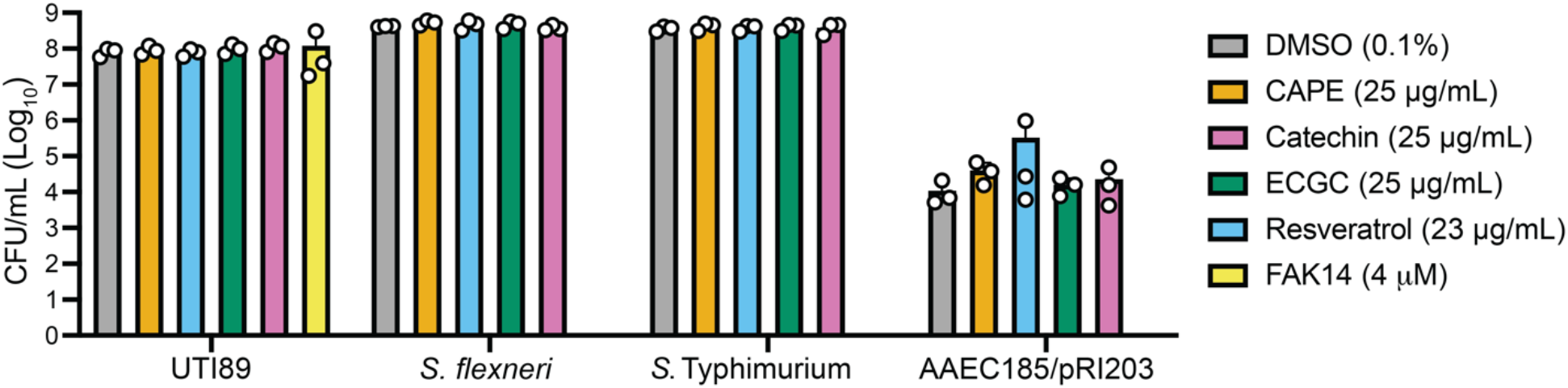
Impact of the phenolic compounds and FAK14 on bacterial viability. Bacteria were inoculated into 24-well plates containing complete RPMI medium ± the indicated drugs or DMSO (carrier alone), as described for the cell association and invasion assays but without host cells present. After 2-h incubations in a humidified CO_2_ incubator at 37°C, bacterial titers were enumerated by plating serial dilutions. Bars indicate mean values from 3 independent assays (with SD).

